# Prediction and control of symmetry breaking in embryoid bodies by environment and signal integration

**DOI:** 10.1101/506543

**Authors:** Naor Sagy, Shaked Slovin, Maya Allalouf, Maayan Pour, Gaya Savyon, Jonathan Boxman, Iftach Nachman

## Abstract

During early embryogenesis, mechanical signals, localized biochemical signals and neighboring cell layers interaction coordinate around anteroposterior axis determination and symmetry breaking. Deciphering their relative roles, which are hard to tease apart in vivo, will enhance our understanding of how these processes are driven. In recent years, in vitro 3D models of early mammalian development, such as embryoid bodies (EBs) and gastruloids, were successful in mimicking various aspects of the early embryo, providing high throughput accessible systems for studying the basic rules shaping cell fate and morphology during embryogenesis. Using Brachyury (Bry), a primitive streak and mesendoderm marker in EBs, we study how contact, biochemical and neighboring cell cues affect the positioning of a primitive streak-like locus, determining the AP axis. We show that a Bry-competent layer must be formed in the EB before Bry expression initiates, and that Bry onset locus selection depends on contact points of the EB with its surrounding. We can maneuver Bry onset to occur at a specific locus, a few loci, or in an isotropic peripheral pattern. By spatially separating contact and biochemical signal sources, we show these two modalities can be integrated by the EB to generate a single Bry locus. Finally, we show Foxa2+ cells are predictive of the future location of Bry onset, demonstrating an earlier symmetry-breaking event. By delineating the temporal signaling pathway dependencies of Bry and Foxa2, we were able to selectively abolish either, or spatially decouple the two cell types during EB differentiation. These findings demonstrate multiple inputs integration during an early developmental process, and may prove valuable in directing in vitro differentiation.

## Introduction

In the early mouse embryo, the blastocyst implants into the uterus wall on the fourth day of gestation. A series of events follows, among which is the formation of an anteroposterior (AP) axis, initiated by the localization of anterior visceral endoderm (AVE) cells at the prospective anterior side and emergence of the primitive streak (PS) at the posterior side [1]. The PS is formed through interaction between BMP, Nodal, FGF and Wnt pathways at the posterior side of the blastocyst, and limited to that side by inhibitory signals originating from the AVE at the anterior side [1, 2]. Mechanical pressure, presumably stemming from the uterus wall, was suggested to be necessary for proper AP axis formation, including the positioning of the AVE and elongation of the egg cylinder [3], though this dependence was later challenged [4]. Another factor shaping PS and AP axis formation is the visceral endoderm layer, characterized at E5.5 by the expression of Foxa2. As mechanical signals, localized biochemical signals and neighboring cell layer contexts all co-occur in a specific reproducible pattern during embryogenesis, it is very hard to dissect their respective roles and dependencies in specifying PS location selection and axis formation.

In-vitro multi-cellular models such as embryoid bodies (EBs), and gastruloids derived from EBs, have been used in recent years to dissect certain aspects of early embryonic development [5-8]. These models are formed by 3D aggregation of pluripotent stem cells. Given proper conditions, these aggregates will differentiate in a partially organized manner, break their radial symmetry and develop an AP axis [8-10]. Interestingly, when aggregated from a small number of cells and under specific signal conditions, EBs form “gastruloids”, displaying not only symmetry breaking but also axis elongation [8, 11]. Germ layer structure was also shown in confined 2D colonies [5, 12, 13]. More recently, it was shown that by mixing mouse ES cells with extra-embryonic cells it is possible to obtain in vitro structures that resemble early stage mouse embryos, both by gene expression profile and morphologically, including dual axis formation [14, 15]. The success of these in-vitro structures in capturing some of the spatiotemporal aspects of early embryogenesis make them valuable model systems for studying basic principles of early development, in addition to strategies of in vitro differentiation and tissue engineering. One of the earliest fate decisions that can be observed in embryonic stem cells is their differentiation to mesendoderm progenitors, which are characteristic of the PS and are marked by Brachyury [16-18]. In 2D patterned colonies, the biased onset pattern of Bry has been associated with colony geometry as well as cell density [19]. In 3D models such as EBs and gastruloids, the polar onset pattern of Bry expression provides one of the first detectable events of symmetry breaking [8-10, 19, 20]. The mechanisms leading to the spatial selection of this pole and axis formation in these 3D models are, however, not clear.

Here we show that Bry expression onset is contact dependent in EBs. While the entire EB outer shell has the potential to express Bry, the expression of this regulator is initiated at the EB contact point with its surrounding surface. By manipulating the EB’s environment we were able to alter the onset of Bry expression to either multiple loci, to a wide contact surface or to an isotropic pattern throughout the outer shell. By creating spatially separate sources of contact and biochemical signals, we demonstrate how the tissue can integrate these two modalities to define a single Bry locus. Remarkably, Foxa2 appears with an eccentric bias towards the future onset site of Bry, evident of an earlier symmetry breaking event, and revealing different dependencies on Wnt signaling.

Our findings provide insight into symmetry breaking mechanisms in 3D tissue models, and primitive streak spatial selection *in utero*, demonstrating how contact and biochemical signals may integrate to drive symmetry breaking and mesendoderm differentiation.

## Results

### Brachyury expression onset location in embryoid bodies is contact dependent

We have previously observed that in EBs differentiated within agarose microwells, Bry expression predominantly starts from a single locus on the EB external shell, and spreads in that layer towards the opposite pole, where a locus is defined as a cluster of Bry+ cells (see Methods) [20]. However, it is unclear whether the spatial selection of the locus is an outcome of stochastic symmetry breaking within the EB, and why Bry onsets solely on the outer shell and not in the volume. We find that in EBs differentiated in agarose microwells, Bry expression onsets from the EB’s contact point with the microwell’s wall or floor (Fig. 1A-E, Supplementary Movie 1-4). Out of 47 EBs imaged, 16 showed Bry onset at the bottom, 12 showed side onset, 14 showed joint side and bottom onset (Fig. 1B) and 5 showed two disparate onset loci. We noted that side-onset or dual-onset EBs are generally larger than bottom onset ones (mean EB radius=120um, σ=16um, vs. mean radius=95um, σ=17um, P=0.001), raising the hypothesis that side onset is driven by contact with the side wall, which is more common in larger EBs. This is reinforced by examining EBs with two Bry loci, which we found to be touching the microwell boundaries at two points or a large surface area overlapping the onset loci (Fig. 1C). We therefore compared in a separate experiment the onset angles in bottom-only contact EBs (n=9) with EBs touching the side wall (n=10), finding significant difference indicating that indeed onset location is affected by contact (Fig. 1E, p=3e-5). Interestingly, EBs differentiated in hanging drops also display a single locus Bry pattern (Fig. 2A,B). Since EBs grown in hanging drops lie at the drop’s bottom boundary, we hypothesized their onset may also be biased toward this boundary. Analyzing the locus location in hanging drops (n=12), we indeed find that Bry locus onsets from the bottom, similarly to the bottom-contact EBs grown in microwells (Fig. 1E). Taken together, we find both mechanical contact and confinement by liquid to air interface can determine the onset location of Bry in EBs. Such contact dependence may be explained by signal consolidation around the contact interface and/or by mechano-sensing pathways.

**Fig 1.**
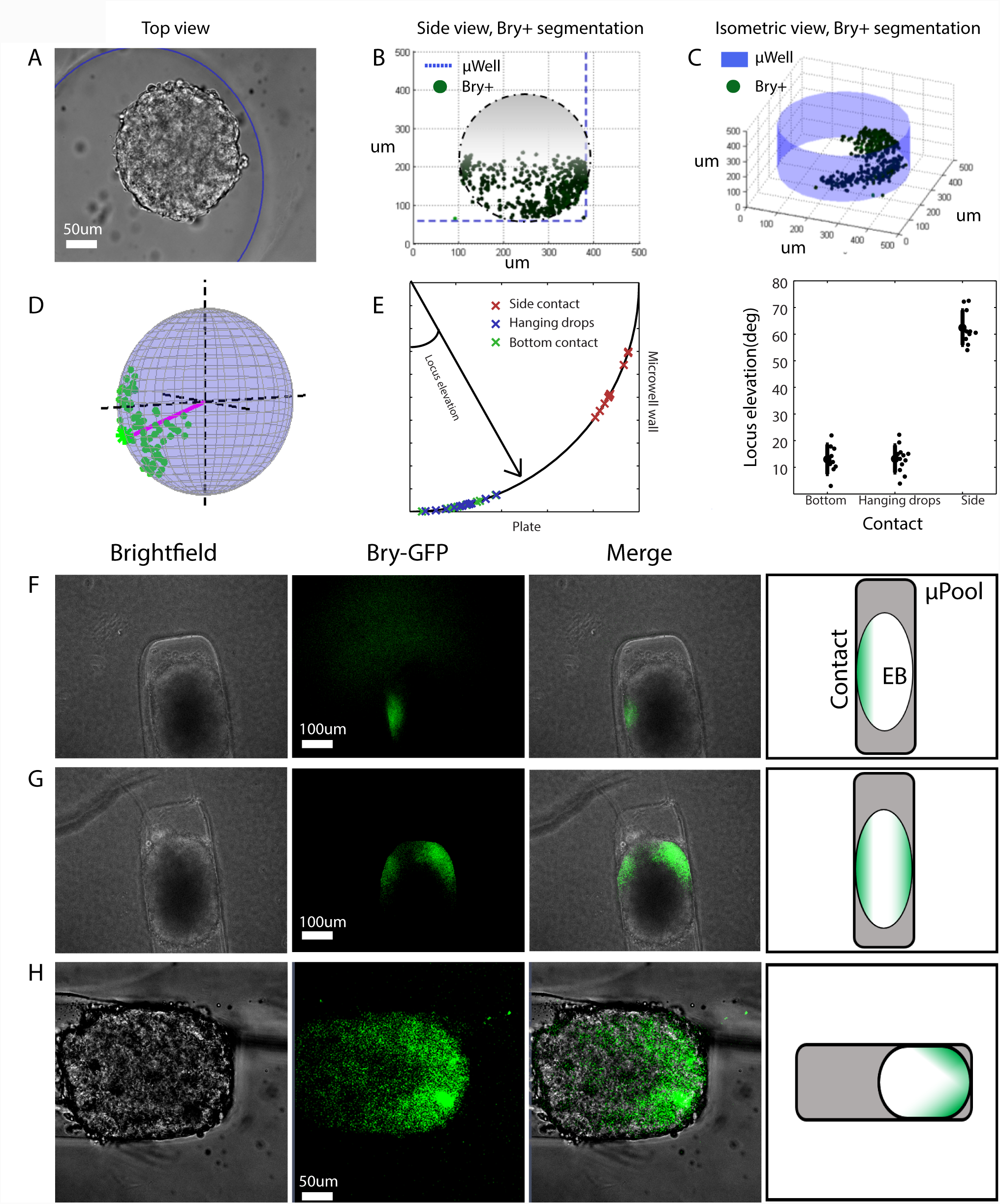
Bry activation is contact dependent. (A) An EB in a microwell at 72 hr from aggregation. Microwell walls are outlined with a blue circle. (B) Segmented Bry signal for the EB from panel A, at sagittal view aspect, showing Bry onset from the contact. (C) An EB touching the well walls at two points. Bry onsets from both points. (D) Estimating the spatial location of Bry locus on segmentation data. The locus vector is shown in magenta. The blue sphere denotes the full EB volume. (E) Locus elevation angle for EBs with bottom contact only (red marks) or side contact (green marks) as determined by brightfield channel at t = 60hr, depicted on a quarter circle (left) or as mean with std error bars (right). Locus elevation is largely determined by contact point. For bottom onset, the locus angle is biased upwards due to asymmetric expansion of Bry+ cells in azimuth. For side onset, the locus angle is biased downwards due to a positive slope of the microwell walls. (F-H) Enforcing contact points by growing EBs in narrow micropools (200 um wide) affects Bry locus locations: Single side locus (F), double side loci (G) or an EB with its right cap fully touching the micropool walls onsetting from the entire cap (H).

**Fig 2.**
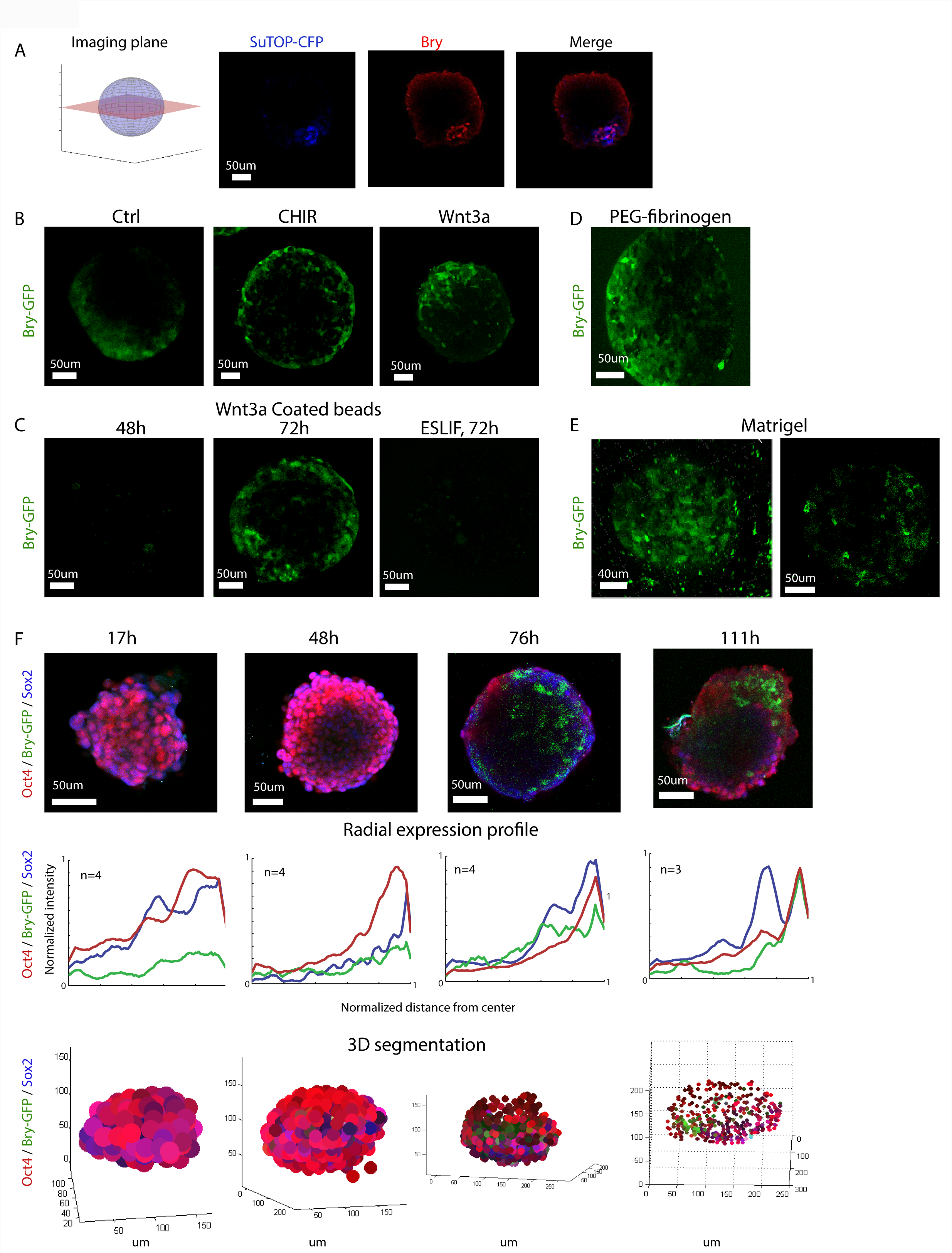
A Bry competent layer is formed prior to onset on the outer shell of the EB. (A) Canonical Wnt activation (measured in EBs aggregated from SuTOP-CFP mES cells in hanging drops) is co-localized with Bry at its onset locus. Left: depiction of the equatorial slice, used throughout the figure. (B) Treatment with CHIR between 0-72 hr (middle) triggers Bry onset on the entire shell, compared to control (left), or soluble Wnt3a perturbation (right), which show a polarized onset. (C) EBs embedded with Wnt3a-coated microbeads at 48 hr (left) and 72 hr (middle) of differentiation, or under pluripotency conditions (right). (D) Embedding the EB in a PEG-fibrinogen hydrogel at 24h has no effect on Bry locus (E) Embedding in Matrigel leads to Bry onset isotropically in most EBs (13/15 EBs, left) or from a single locus in some large EBs (2/15 EBs, right). (F) Immunostaining of Sox2 and Oct4 in E14 Bry-GFP EBs at the indicated time points after aggregation. Top row: equatorial slices. Bottom row: representations from whole-EB image-stack 3D segmentation, showing Oct4 as red level, Sox2 (blue level) and Brachyury-GFP (green level). Oct4 recedes towards the perimeter by 48 hours. Middle: normalized radial intensity profiles of Oct4, Bry and Sox2, computed in the direction of maximal expression on the outer shell, averaged over the indicated number of EBs.

We next wanted to check whether the timing and location of Bry onset can be controlled by imposing contact at a specific locus at an early stage. To this end, we fabricated 200um wide rectangular “micropool” structures to enforce early contact at two sides of the EB (Fig. 1F). EBs grown in these micropools expressed Bry at their contact points with the micropool walls (13/14; Supplementary Movie 5, 6). Out of these, 11/14 showed a single locus onset (Fig. 1F), 2/14 showed simultaneous onset at two loci (Fig. 1G), and in one instance, when positioned at the end of the micropool, at the entire contacting cap (Fig. 1H). EBs grown in wider micropools (500um width) and which did not touch the walls showed bottom Bry expression onset, at the contact point with the plate floor (4/5), similar to the case in large microwells (n>50 in multiple experiments [20]). We then evaluated the effect of early contact on the timing of Bry onset. When grown in the narrow (200um) micropools, EBs created contact with the micropool sides earlier (50-60 hours), but Bry onset did not occur earlier compared to the non-touching controls in the wide micropools (79h, σ=2 hrs, n=11 vs. 75h, σ=5 hrs, n=5). This suggests that cells on the outer shell of the EB did not yet have the potential to express Bry when contact was initially applied to the EB, and more molecular changes had to take place before Bry onset could occur around the contact point.

### Brachyury competent shell is formed prior to onset

The observation that EBs can form multiple contact loci or an entire cap contact locus when confined in micropools raised the question whether the potential to express Bry is limited to specific areas on the outer layer of the EB, to the entire shell, or is uniform throughout the EB volume. Growing EBs in hanging drops, we first confirmed that a canonical Wnt activation reporter (SuTOP-CFP, [21]) showed co-localization with Bry, in agreement with observations that Bry is activated via the canonical Wnt pathway (Fig. 2A) [9, 22]. We then tested EB spatial limitations for Bry onset under external Wnt induction. To this end, we induced the canonical Wnt pathway by either adding CHIR99021 (CHIR), a small-molecule Wnt pathway agonist bypassing the pathway receptor via Gsk3β inhibition, by adding soluble Wnt3a ligand, or by embedding Wnt3a-coated beads within the entire EB volume during aggregation (Fig. S1A) [23]. EBs under CHIR treatment showed isotropic onset, with no distinct locus, suggesting that the entire outer shell is Bry competent (n>10, Fig. 2B). Activation by soluble Wnt3a resulted in a stronger, yet polarized locus (Fig. 2B, right). EBs embedded with Wnt3a beads from aggregation displayed isotropic or near isotropic Bry expression onset on the outer shell at 72 hours, similar to CHIR treated EBs, which can be attributed to multiple loci caused by different beads. As in our forced early contact results, Wnt3a beads did not lead to Bry expression up until 72hrs after aggregating the EBs, nor did they induce Bry under pluripotency conditions (Fig. 2C). These results confirm that similar to their susceptibility to contact, outer-layer cells need to reach a certain maturation stage before becoming Wnt-susceptible to activate Brachyury.

As the entire outer shell is Bry-competent, and as contact is associated with Bry onset locus, we hypothesized that given uniform contact conditions, outer layer cells may induce Bry in a spherically isotropic pattern. We therefore tested the Bry onset pattern when EBs were differentiated while embedded in hydrogels simulating either poor (PEG-fibrinogen) or rich (Matrigel) ECM composition. When embedded in PEG-fibrinogen hydrogel, the EB kept a single locus onset despite having an isotropic contact with the gel (Fig. 2D). When embedded in Matrigel, EBs in a range of sizes displayed isotropic onset pattern (13/15, mean radius at onset=95, σ=50um), while two large EBs (r=120um, 180um) initiated Bry from a distinct locus (Fig. 2E, Supplementary Movie 7, 8). When differentiating uniformly small EBs in microwells (mean EB radius at onset=78um, σ=10um, n=9), Bry-GFP showed a uniform isotropic onset (7/8 EBs, average onset time 78h, σ=9h), similar to small EBs embedded in Matrigel. This suggests an isotropic pattern can be a result of size, which may be explained by a minimal locus size that is larger than the small EB surface (Fig. S1B), or a minimal signal gradient requirement for a polarized pattern. In summary, given sufficient EB size, a polarized Bry pattern can be obtained at contact points, but also under uniform gel contact, suggesting this is a robust EB behavior. However, as our results from CHIR induction, Wnt3a bead embedding and growth in Matrigel demonstrate, the entire outer shell is competent to activate Bry, given strong activation of the Wnt pathway.

Previously, Oct4 and Sox2 expression levels in differentiating ES cells were suggested to direct their differentiation toward mesendoderm or neural ectoderm, respectively [24]. To test whether the competence of the outer shell can be explained by the expression pattern of these early pluripotency factors, we immunostained for Oct4 and Sox2 at 17, 48, 76 and 111 hrs into differentiation. We indeed observed that the expression of both genes recedes from the center towards the outer shell with time, and by 72 hrs both Sox2 and Oct4 are highly expressed only at the outer shell of the EB (Fig 2F).

Which molecular components may mediate the effect of contact on Bry onset in outer layer cells? Previously, fibronectin was shown to induce Bry expression, and a mechanism linking fibronectin to Wnt signaling was proposed [25]. We therefore hypothesized such a link can explain both the relation between contact and Wnt signaling (leading to Bry onset), and the location choice for the Bry locus. We found fibronectin is expressed isotropically on the EB’s outer shell, implying that its mere expression does not determine the location of Bry onset (Fig. 3A). We then inhibited fibronectin activity using an anti-Fn1 antibody, starting at 0, 24, 48, or 72 hours following aggregation. While fibronectin inhibition at aggregation or 24hr inhibited Bry expression, it had little to no effect when added at 72hr and a mixed effect was identified when added at 48hr (Fig. 3B). This suggests that the dependence of Bry positive fate on fibronectin is limited to early stages, while the accumulation of Brachyury in the cells, as well as the expansion of its expression to neighboring cells do not depend on fibronectin.

**Fig 3.**
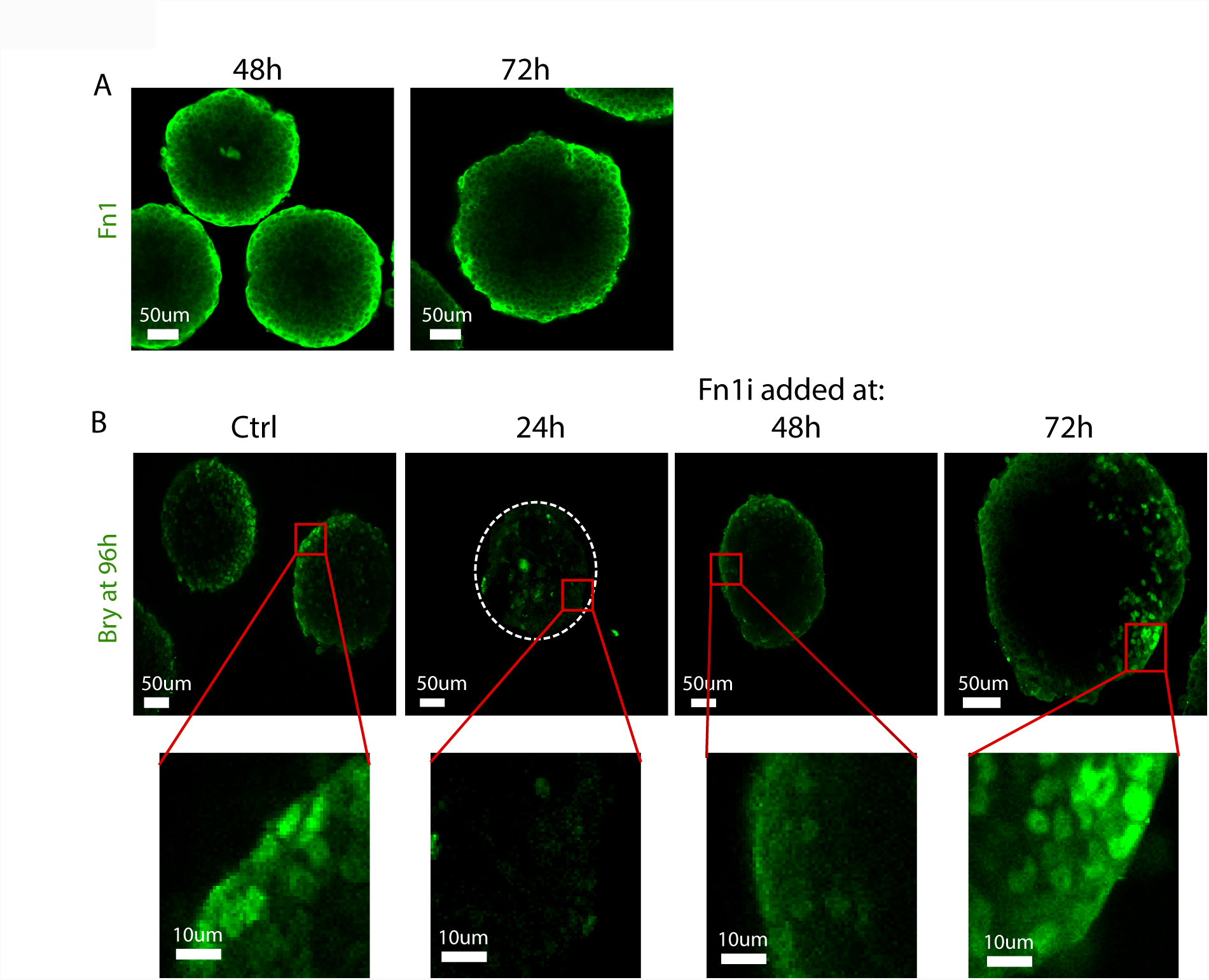
Fibronectin is expressed isotropically on the EB outer shell, and is needed at early stages for Bry expression. (A) Fibronectin immunostaining at 48 and 72 hr from aggregation. Fibronectin is expressed uniformly on the outer shell at least until Bry onset. (B) EBs subjected to fibronectin inhibition by an anti-Fn1 antibody starting at 24, 48 or 72 hr from aggregation, and immunostained for Bry (green) at 96 hr. Fibronectin inhibition prevents Bry onset, but only when started earlier than 48 hours.

### Wnt and contact signals are integrated to activate Bry

Our results suggested that both Wnt signaling and mechanical contact affect Bry expression onset, with no clear functional hierarchy between them. To test how these two signal modalities relate to each other in this context, we designed a system where we create a spatial separation between surface contact point and a local source of Wnt signaling. To this end, we transfected HEK cells with a plasmid constitutively expressing either Wnt3a-P2A-H2B-CFP, Dkk1-P2A-H2B-Venus or, for the control group, H2B-CFP (Fig. 4A). A clump of HEK cells was then integrated into each EB at a random location shortly after aggregation. The EBs were then differentiated, as in the wild-type experiments, and the location of Bry locus was analyzed with respect to the contact point and the locus of signaling cells using two-photon microscopy (Fig. 4B, C, D). In the EBs containing Dkk1 or Wnt3a cell clumps, the Brachyury locus overlapped with the plane defined by EB centroid, HEKs, and contact point (n=16; average deviation from co-planarity 9.3 degrees) (Fig. S2), which allowed us to analyze the joint effects of contact and Wnt3a/Dkk1 signaling on Bry expression locus as angular deviations on a single plane (Fig. 4B). The amount of Wnt3a secreting agents determined the effect on Bry locus bias from the contact. For small clumps of Wnt3a expressing cells (3-5 cells), the source of Wnt3a leads to a shift in the Bry locus, so that it is positioned between the contact point and the Wnt3a expressing locus (Fig. 4C, E-G). For larger clumps of Wnt3a secreting cells (>10 cells), the Bry locus was pulled closer to the HEKs cluster (Fig. 4E, F). In contrast, a source of Dkk1 shifts Bry locus away from the contact point (Fig 4D, E, G). This indicates that Dkk1 can exert its limiting effect over at least 150um. Moreover, the locus shift from the contact point, towards or away from the signaling source shows how Wnt3a and Dkk1 signal integration determines the locus location. In control EBs (harboring clumps of H2B-CFP cells) we see no spatial effect on the locus of Bry onset, which is mostly at the contact point (3/4). These results suggest that contact and Wnt effects on Bry onset are integrated rather than override each other. One possible explanation is that contact mediates Wnt signaling (which then gets spatially summed with the other source of Wnt signal). It is also possible, however, that contact acts through a different signaling pathway, which gets summed with the Wnt pathway in triggering Bry onset.

**Fig 4.**
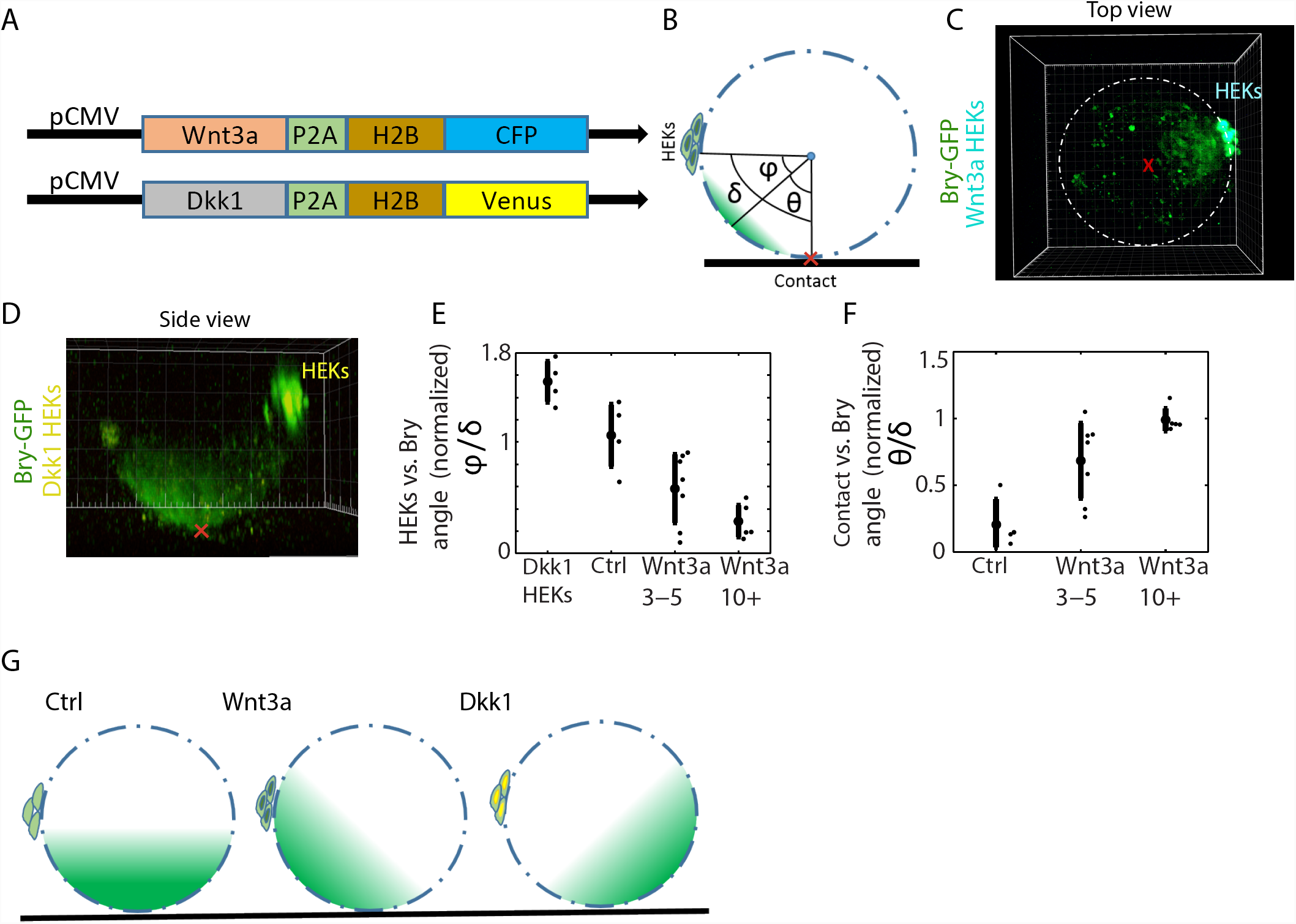
Bry onset locus is determined by contact and Wnt signaling integration. (A) Wnt3a and Dkk1 constructs used for ligand secretion from HEK cells (B) A signal source (a clump of transfected HEK cells) is embedded into each EB. The pairwise spatial angles between signal source, contact point with well bottom surface (red x mark) and Bry locus are computed. (C) Example of an EB with a Wnt3a signaling source. Bry onset locus is biased towards the Wnt3a secreting HEKs (D) Example of an EB with a Dkk1 signaling source. Bry onset locus is biased away from the Dkk1 secreting HEKs. (E) Angle between HEK signaling source and Bry locus (φ) normalized by the angle between the HEKs and the contact point (δ). Wnt3a secretion biases onset from the contact point towards the Wnt source in a quantity dependent manner, where Dkk1 secretion biases the locus away to the opposite side. (F) Angle between contact point and Bry locus (θ) normalized by the angle between the HEKs and the contact point (δ). Bry is pulled further away from the contact as the number of cells in the Wnt3a secreting HEKs clump grows. (G) A schematic description of the spatial bias effect.

### Foxa2 is a precursor to symmetry breaking and Bry onset locus

To explain the choice of Bry onset location in the absence of external contact surfaces (i.e. in hanging drops), we examined additional genes that are predicted to be expressed at or near the primitive streak before gastrulation. We have previously suggested that once Brachyury onsets, the EB progresses at a common developmental pace, delineating a differentiation plan associated with multiple early development genes [20]. We therefore hypothesized that the spatial expression pattern of key proteins in EBs before and during Bry onset will show similarity to their *in utero* pattern around the primitive streak. To test this, we immunostained EBs differentiated in hanging drops for several early markers at different time points from aggregation, and checked how their spatial expression patterns relate to that of Bry. We find that Eomes, a primitive streak marker regulated by Nodal and an EMT driver, was localized to the same pole as Bry though expanding further on the outer shell (Fig. 5A). E-cadherin was expressed throughout the EB prior to Bry expression onset, and then its expression declined where Bry was expressed, indicating that epithelial mesenchymal transition takes place in the EB (Fig. 5B). Loss of E-cadherin enables cell movement, which is supportive of previous observations of gastrulation-like events in EBs and 2D models [5, 8, 20], and is consistent with primitive streak behavior. Oct4, whose expression initially declines in the inner layers, is gradually receding on the outer shell towards the opposite pole as Bry expression expands, similar to the opposing gradients observed in late streak stage epiblast (Fig. 2F, 5C).

**Fig 5.**
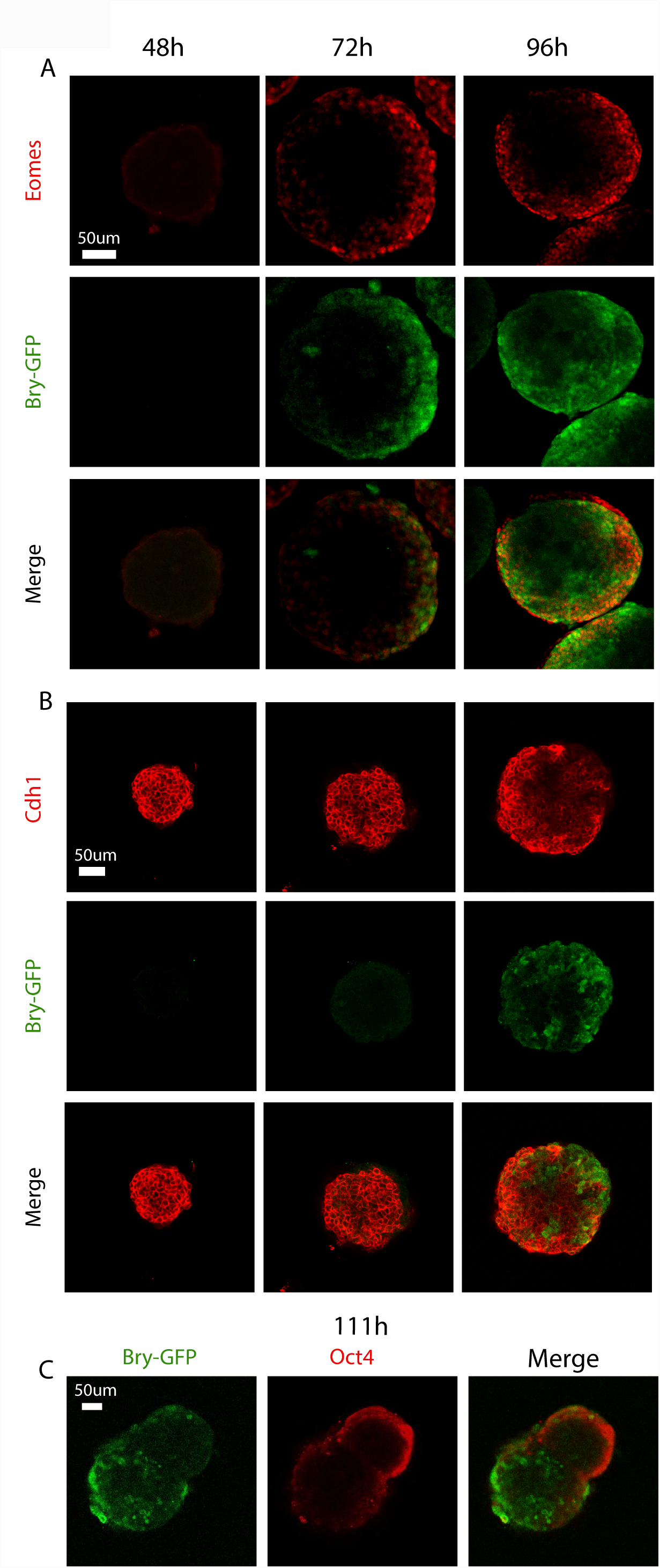
Brachyury onsets in a primitive streak like area in EBs. Immunostaining of several early developmental markers, comparing their spatial expression patterns to that of Brachyury. (A) Eomes onsets with Bry at the same locus and extends beyond Bry spatial expression (B) E-cadherin recedes from the locus where Bry appears, allowing cell motility (C) Oct4 is expressed on the outer shell, then receding to the pole opposite that of Bry onset.

The examples above demonstrate how the expression and spatial arrangement of different proteins in the EB during pre-streak through streak phases resemble the equivalent in vivo stages, albeit with an altered topology. Since in the embryo the primitive streak is initiated adjacent to the visceral endoderm layer, we tested for spatio-temporal correlation of Foxa2, a visceral endoderm marker, with Bry in the EB. Such a correlation could point to an additional driving factor in Bry locus determination. Indeed, Foxa2 expression appeared up to 24 hours prior to Bry at an internal area within the EBs, with Bry onset following on the shell juxtaposed to Foxa2 expressing cells (Fig. 6A), suggesting a common driver or inter-layer dependency. This spatial relation between Foxa2 and Bry is accentuated in large EBs (∼700uM in diameter at onset) (Fig. 6A, bottom, Fig. S3A), and suggests these two cell types have a regulatory relation.

**Fig 6.**
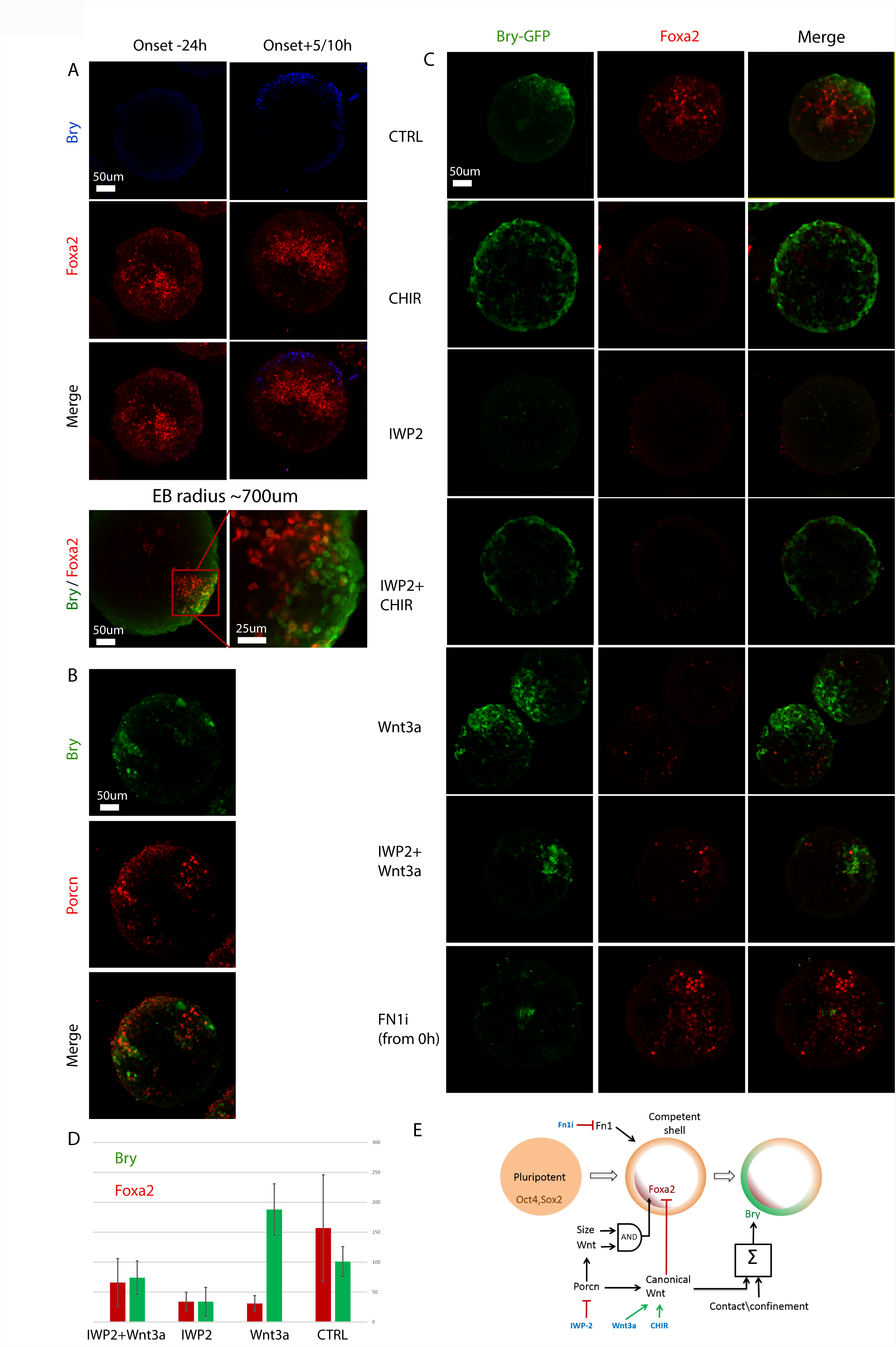
Foxa2 is a Wnt dependent precursor to symmetry breaking and Bry expression locus. (A) Foxa2 expression is juxtaposed to Bry+ cells in EBs, appearing >24 hr before the onset of Bry in multiple experiments (n>20). Bottom: Foxa2/Bry pattern in a large (700um) EBs (similar pattern seen in 3/3 large EBs). (B) Porcn expression partially overlaps with Bry, suggesting that Wnt ligands can be secreted in these cells. (C) CHIR upregulates Bry, triggers an isotropic expression pattern on the Bry-competent outer shell while downregulating Foxa2 expression in the EB; IWP2 (a porcupine inhibitor) annihilates Wnt secretion, downregulating Foxa2 and Bry; CHIR rescues Bry under IWP2 perturbation, however Foxa2 is still downregulated, suggesting that it is not dependent on canonical Wnt; Wnt3a up regulates Bry while down regulating Foxa2; Wnt3a partially rescues both Bry and Foxa2 under IWP2; Fn1 inhibition abolishes Bry expression however leaves Foxa2 intact. All signals were applied from 48 hr after aggregation, except Fn1 inhibition, applied at 0 hr. (D) Perturbation effects on Bry and Foxa2 expression quantified as number of highly expressing cells per EB. (E) A schematic model summarizing the factors affecting Bry onset in the EB.

To look for candidate genes that may convey the regulation of Bry next to Foxa2+ cells, we analyzed a published single cell expression dataset from E6.5 mouse embryos [26]. We found that within epiblast cells, both Foxa2 expressing cells and Bry expressing cells are significantly enriched for Porcupine (Porcn) expression (p=2.6e-8 and p=8.83e-10, respectively). Porcupine palmitoylation of Wnt ligands is required for their secretion, suggesting that Foxa2 positive cells in EBs may secrete Wnt ligands to the adjacent perimeter, thus promoting the onset of Bry in an adjacent locus. We therefore immunostained for Porcn in differentiating EBs, finding that porcupine expression is spatially limited to areas adjacent to and overlapping Bry onset (Fig. 6B). Inhibiting Porcn activity with IWP-2 starting either at aggregation or 48hr after aggregation (when Foxa2 is already expressed in the volume) resulted in the abolishment of both Bry and Foxa2 (Fig. 6C; Fig S4A). This suggests that both genes require Wnt signaling for their expression. When introducing IWP-2 with CHIR to the EBs 48hr after aggregation, we were able to rescue Bry, however Foxa2 was fully downregulated. When Wnt3a was added to IWP-2 we obtained a partial rescue of both Bry and Foxa2. Interestingly, when perturbing the EBs with just CHIR or Wnt3a, we obtained a similar result where Bry is over-expressed and Foxa2 is either fully (in the case of CHIR) or partially downregulated and spatially decoupled from Bry (in the case of Wnt3a) (Fig. 6C, D, Fig. S3B). These results suggest that an intricate, likely dynamic, signaling balance involving canonical Wnt determines and maintains the fate of these cells, where non-canonical Wnt pathways may be involved in activating Foxa2. In small EBs (<65um radius) Foxa2 had small to negligible expression while Bry encompassed the entire EB (5/5, Fig S4B). This suggests that the EB requires a certain volume (or lengthscale) to allow the development of both mesendoderm and visceral endoderm tissues, consistent with results in 2D colonies [5, 13]. The threshold size may be required to establish sufficient differences, or gradients, in signaling activity. Finally, while Bry expression was abolished under fibronectin inhibition starting at aggregation, Foxa2 showed wild type expression, pointing to a fate decision made at least 24 hours before Bry onset and leading to Foxa2+ cells (Fig. 6C, bottom).

Together these results show how Bry and Foxa2 are spatially related under Wnt signaling. Bry is monotonically upregulated by canonical Wnt where Foxa2 is showing a more complex regulation nature, where it is upregulated by Wnt but downregulated by strong canonical Wnt activation (Fig. 6E).

## Discussion

Here we dissect the factors and mechanisms affecting the polarized expression pattern of Brachyury expression in embryoid bodies. These patterns can be formed in EBs and gastruloids with no extra-embryonic tissue and no external Nodal or BMP sources [8, 11, 20]. We show that physical contact with surfaces, a localized source of Wnt signaling, and proximity to cells expressing genes that characterize the visceral endoderm, may all induce or predict the onset of Brachyury. Interestingly, in the embryo, all these three conditions co-occur near the future location of the primitive streak, making it hard to dissect their respective roles. Brachyury onsets during gastrulation near a source of Wnt signaling (arising from a positive feedback loop between Bmp4 sourced from the ExE and Nodal and Wnt at the posterior epiblast), adjacent to the visceral endoderm layer, in a region that may experience high mechanical pressure due to implantation. Here we analyze some of the hierarchy between these three PS predictors (Fig. 6E).

Brachyury onset polarization is robust to different external conditions and EB parameters, suggesting it is the “default” behavior in this system. A similar result was demonstrated in mouse embryos grown in-vitro, where while mechanical contact may direct AP axis formation and positioning of the AVE, these can also robustly form in hanging drops [3, 4]. Breaking the polar pattern in EBs requires specific intervention, such as continuous canonical Wnt activation by CHIR or applying a few contact points. Contact with external surfaces, or with the hanging drop boundary surface, breaks the EB symmetry and strongly biases the location of Bry onset. We show how through controlled contact we can generate two Bry loci, or a wide “cap” locus, depending on the geometry of the surrounding surfaces (Fig. 1G, H). The fact that Bry onsets in hanging drops or large wells similarly to the way it does in narrow micropools, suggests it is not the effect of mechanical pressure, but rather the interface with contact surface or geometrical confinement. These could act through limiting the diffusion of secreted signal molecules, such as Wnt ligands, resulting in locally higher signal concentration [27]. Another option is bypassing the ligand activation mechanism through interfacing with outside surfaces by integrin-mediated signaling [25], although the localization of Bry locus in hanging drops to the bottom interface suggests that diffusion confinement may be sufficient. Interestingly, the factors we find to affect Bry locus in 3D are very different from the case of 2D patterned colonies, where a recent study related the percentage of Bry+ cells to cell density, and their localization to geometrical confinement or high curvature at the tip of elongated colonies [19].

We find that we can “pull” or “push” the Bry locus away from the contact point using an external source of Wnt3a or Dkk1 secreting cells, respectively. This demonstrates that contact and biochemical signal modalities can be summed by the EB, resulting in a single locus, as opposed to the two loci, or one locus overriding the other we observe in the forced contact experiments. This can be attributed to the gradient strength or to the surface distance between the loci, which may be too large for signal integration to have an effect. One possible mechanism for this integration can be activation of Wnt signaling through the mechanical contact. The dependence of Bry activation on fibronectin may point to such a mechanism [25]. Interestingly, the mechanism proposed by Cheng et al. [25] bypasses the Wnt pathway receptors, and therefore cannot explain why Dkk1 diverts Bry onset away from the contact point, suggesting other mechanisms may be at play. Another option is that Wnt ligands consolidate at the contact surfaces, or have limited diffusion range near them, leading to locally higher concentration [27]. The isotropic peripheral onset we observe in small size EBs may be attributed to a lack of Wnt gradient formation.

The relation of Bry onset to Foxa2 positive cells provides some interesting insights. First, from the differential response to fibronectin inhibition, we show that the decision junction of differentiating ES cells within the EB toward Bry and Foxa2 tissue layers takes place at 24-48h prior to Bry onset. The early appearance of Foxa2 (compared to Bry) suggests these cells, at least at that stage, represent visceral endoderm rather than primitive streak or definitive endoderm [28]. Second, the observation that small-size EBs do not express Foxa2 suggests that a signal gradient may be controlling layers segregation between mesendoderm progenitor and visceral endoderm. This is reinforced by the complex dependency of Foxa2 expression on Wnt signaling, as well as by the close spatial dependency we see between these two cell types in large EBs (Fig. 6, S3). The fact that different perturbations can lead to Bry or Foxa2 expression while abolishing the other fate in the EB, and that the spatial adjacency of these two cell types can be decoupled using external Wnt signaling, suggests the observed spatial relation may be due to both proteins depending on Wnt signaling, though in a different manner.

In contrast to its location, the timing of Bry expression onset in EBs is not affected by external or localized Wnt3a signaling, or by inducing early contact in narrow microwells. These results point to the maturation needed in outer shell cells before they become Wnt and/or contact-responsive to induce Bry. Further characterization of the chromatin or expression changes defining that “mature” state are needed.

In summary, through different perturbations on a 3D in vitro system we were able to delineate basic dependencies of an early fate decision on contact and biochemical factors that co-occur in the embryo. The insights we gained here improve our understanding on the forces shaping differentiation within in vitro systems such as EBs and gastruloids, and shed more light on how they are integrated in vivo.

## Methods

### Cell culture

E14 Bry-GFP mouse ESCs (kindly provided by Dr. Gordon Keller), R1 mouse ESCs and SuTOP-CFP,AR8-RFP mouse ESCs (kindly provided by Dr. Palle Serup) were cultured on gelatin surface using standard conditions on irradiated primary mouse embryonic fibroblasts and knockout DMEM containing 15% fetal bovine serum, 50 ug/ml penicillin/streptomycin, 2 mM L Glutamine, 100 μM non-essential amino acids, 80 μM ß-mercaptoethanol and 10^3^ U/mL LIF.

### EB formation and differentiation

Embryoid bodies were formed in hanging drops. Each drop contained 25ul differentiation medium (IMEM containing 20% FBS, 50 ug/ml penicillin/streptomycin, 2 mM L Glutamine, 100 μM non-essential amino acids, 100 μM ß-mercaptoethanol) with approximately 300 mES cells (100 cells for small EBs, 2000 cells for large EBs). Cells were aggregated for 24\48 hours before being transferred to micropools/ microwells for imaging in the same medium. Alternatively, the EBs were kept in hanging drops for the entire duration of experiment before being imaged or fixed and immunostained. When adding inhibitors or activators to hanging drops we aggregated the EBs in 20ul hanging drops and then added at the appropriate time 5ul of medium with the required substance. For signal perturbation experiments, differentiation medium was supplemented with CHIR99021 (5.35 uM) or recombinant Wnt3a (500 ng/ml). For fibronectin inhibition, the medium was supplemented with anti-Fn1 antibody (Abcam ab6328). For incorporating Wnt3a-bound beads within the EB, prepared beads (kindly provided by Shukri Habib) were pre-mixed with mES cells before distributing the suspension to 25ul hanging drops for aggregation. We added 12K beads to 10ml of medium with the mES cells, averaging at 30 beads per EB. Beads integration into the EBs was confirmed by two-photon microscopy (Fig. S1A).

### Hydrogel and Matrigel

PEG DA fibrinogen was mixed with Igracure 2959 photoinitiator dissolved in EtOH in a 99:1 ratio. The Hydrogel was polymerized under a 365nm lamp for 10 minutes on a glass surface to allow two photon imaging. 24 hours after aggregation, EBs were individually transferred from hanging drops to the gel with a 10ul micropipette prior to polymerization and grown in Iscove medium. Matrigel (Gibco Geltrex A1413202) was thawed on ice in 4 degrees overnight. Matrigel drops (20ul each) were deposited on the glass bottom of a 24-well. The EBs were transferred to into the drops at 48h after aggregation, and after gel solidification the wells were filled with medium.

### HEK incorporation in EBs

HEK 293 cells were split to three 12-wells (50% confluence). 12 hours later, the wells were transfected with constitutive pCMV-H2B-Cerulean, pCMV-Wnt3A-P2A-H2B-Cerulean or pCMV-DKK1-P2A-H2BVenus plasmids, using the Xfect™ transfection reagent (Cat.# 631318). 24 hours post transfection, the transfected HEKs were trypsinized, suspended as single cells and injected into hanging drops containing 12-hour old E14 Bry-GFP EBs at different concentrations – 10/20/50 HEKs per EB. 24 hours later, the injected EBs were transferred into microwells and imaged using a two-photon microscope.

### Micropools patterning with Deep Reactive-Ion Etching (DRIE) and microwells

To create the silicon template for the shaped micropools and microwells we have implemented a high aspect ratio etching system usually used for microelectronics fabrication. With DRIE we were able to determine the micro pattern’s exact depth by selecting a wafer with a specific handle/ device thickness. This suggests an advantage over the common photoresist techniques, which requires fine-tuning of the process to have an exact pattern depth. We have fabricated 200-500um diameter microwells and 100-500um wide micropools, with a depth of approximately 400um. The patterns were then coated with PTFE to minimize friction with the template when pulling the elastomer out. To create the microwell/pool negative, PDMS elastomer (Sylgard 184) mixed with a curing agent (1:10) was poured into the silicon wafer mold and incubated o/n at 50C. Individual stamps were then cut out using a scalpel. To generate the positive agar microwells/pools we placed the stamps face down inside poly-D-Lysine coated glass bottom plates and injected agarose under the stamps. The plates with the injected stamps were then placed in a vacuum chamber for 90 minutes, followed by another round of agar injection and additional 10 minutes in the vacuum chamber. The stamps were then removed, leaving the micro patterns imprinted in the agar.

### Hanging drops imaging

EBs were aggregated in hanging drops on plastics slides. The slides were mounted in the two-photon microscope and imaged with EC Plan-Neofluar 20x/0,50 M27 lens, providing a working distance of 2mm. The data was then segmented in Imaris and analyzed in MATLAB.

### Live imaging

We have sourced data from both 2D and 3D live imaging experiments. 2D epi-fluorescence experiments offer a short acquisition cycle for a large cohort of EBs, thus yielding accurate temporal tracking and statistics based on a large number of samples. On the other hand they do not possess elevation distinction and the superior resolution and cell separation that more advanced microscopy techniques offer. For 3D imaging we used a two-photon microscope with which we were able to have a full view of the signal’s dynamics and track individual cells. However, this technique requires large data analysis, had a longer acquisition cycle, and is less compatible with multiple EB imaging. Two-photon imaging was done using a Zeiss LSM7 inverted two-photon microscope with a 20X/0.8NA air objective. Each EB was scanned at 3um intervals along the *z* direction. Horizontal resolution was set to 512×512 pixels at approximately 0.6um per pixel. GFP was excited at 930nm. RFP, Alexa 488, Alexa 405 and Alexa 594 were excited at 800nm. CFP was excited at 860 nm. Epifluorescence imaging was done using a Nikon TiE epi-fluorescence microscope equipped with a motorized XY stage (Prior) at 10x magnification using NIS Elements software. Acquisitions were taken every 25/30 minutes for 14-72 hours. For live imaging, in both systems we used an Okolab incubation cage, maintaining 5% CO2 and 37C.

### Immunohistochemistry

We fixed the embryoid bodies with 4% Paraformaldehyde o/n at 4C, followed by rinse cycle and an o/n wash in PBS. We add primary antibody (Eomes: Abcam ab23345; Bry: R&D Systems AF2085; Foxa2: Abcam ab108422-100; E-cadherin: Cell Signaling 24E10; Oct4: Santa Cruz sc-8628; Sox2: Santa Cruz sc-365964) with blocking buffer (PBS, 0.1% Triton x100, 5% FBS) and incubate o/n at 4C, followed by four wash cycles in blocking buffer for at least half an hour per cycle. We then add the secondary antibody in blocking buffer and incubate it o/n, followed by four wash cycle as described for the primary antibody. Antibodies concentrations used are the same as recommended by the manufacturer for 2D staining.

### Segmentation and Analysis

The Two-photon microscopy data was contrast enhanced in Fiji and then spot segmented in Imaris, where the integration radius was set to 7um and the filter was set to sum intensity with a threshold of 110/255. The background is subtracted prior to segmentation. The segmented data was read and analyzed in MATLAB. For Bry locus estimation in hanging drops or microwell experiments, locus elevation and azimuth angles were calculated by the sum vector of all Bry+ cells. The EB center and radius was evaluated by Bry+ cells curvature and the angle between the EB bottom point and Bry locus was computed for the HEKs incorporation experiments, we developed a script that calculates geometrical properties between HEKs cluster, Bry locus and contact point. The script detects HEK cells in each time point by channel and intensity. We then calculate the geometrical mean of the HEKs and set it as the HEKs location point. Next we look for Bry locus, where a locus is defined as a minimum of 5 cells above intensity threshold of 100 within a distance of 50um from each other, all in the green channel. Next we define the contact point of the EB with the well bottom as the bottom point of the sphere and calculate the geometric properties between all pairwise combinations of HEKs cluster, Bry locus, and contact point. The geometrical properties calculated are the spatial angles, Euclidean distances and geodesic distances.

## Acknowledgements

We thank Shukri Habib for providing Wnt3a coated beads, Palle Serup for the SuTOP-CFP mES cell line, Dror Seliktar for providing PEG-fibrinogen gel, and David Sprinzak and Omri Wurzel for providing helpful feedback on the manuscript. This study was supported by the Israel Science Foundation (grant No. 1665/16) and by a fellowship from the Edmond J. Safra Center for Bioinformatics at Tel-Aviv University.

---

**Fig S1.**
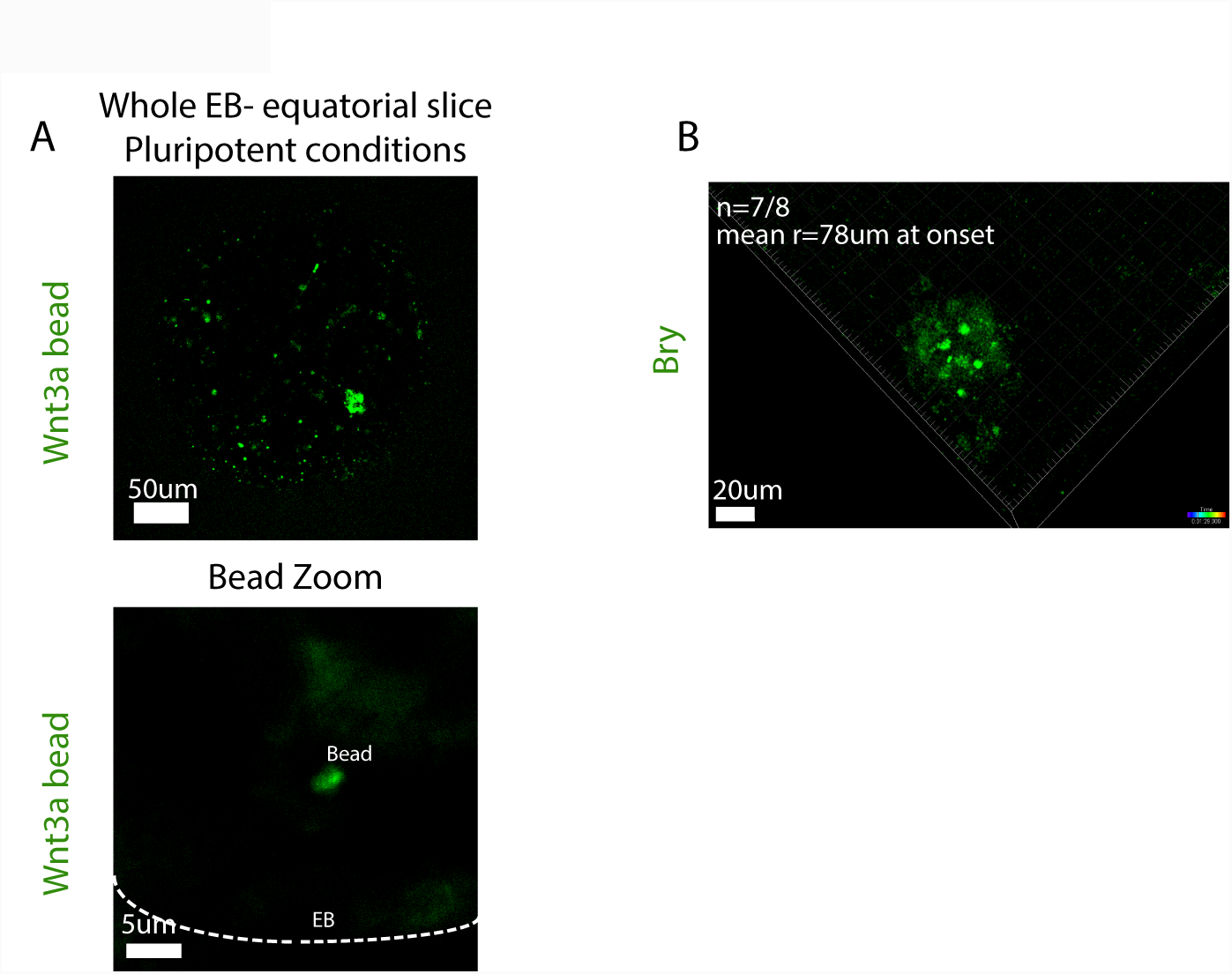
Wnt3a beads integration and small-EB isotropic onset. (A) Top: An equatorial slice of an EB embedded with multiple Wnt3a-coated beads (green). Bottom: A zoom in snapshot on a Wnt3a-coated bead. (B) An example small EB differentiated in microwells at 72 hr from aggregation, showing isotropic (spatially uniform) Bry expression onset. The isotropic pattern occurred in 7/8 small EBs (radius at onset = 78+-10um).

**Fig S2.**
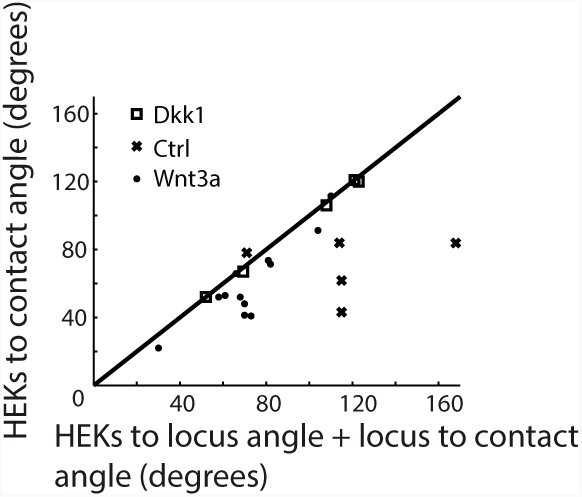
Wnt3a/ Dkk1 source, Bry locus, and Contact point show high co-planarity. The angle between HEKs signal source and contact (δ, Y-axis) vs. the sum of the angle between HEKs to Bry locus and the angle between Bry locus and contact (θ+φ, X-axis) for all EBs quantified in Fig. 3E,F. For EBs harboring Dkk1 or Wnt3a producing HEKs, δ is approximately equal to θ+φ, which indicates that Bry locus inhabits the same plane defined by the HEKs, contact point and the EB centroid. As expected, in control EBs there is a larger deviation from co-planarity, as the Bry locus is not constrained to that same plane.

**Fig S3.**
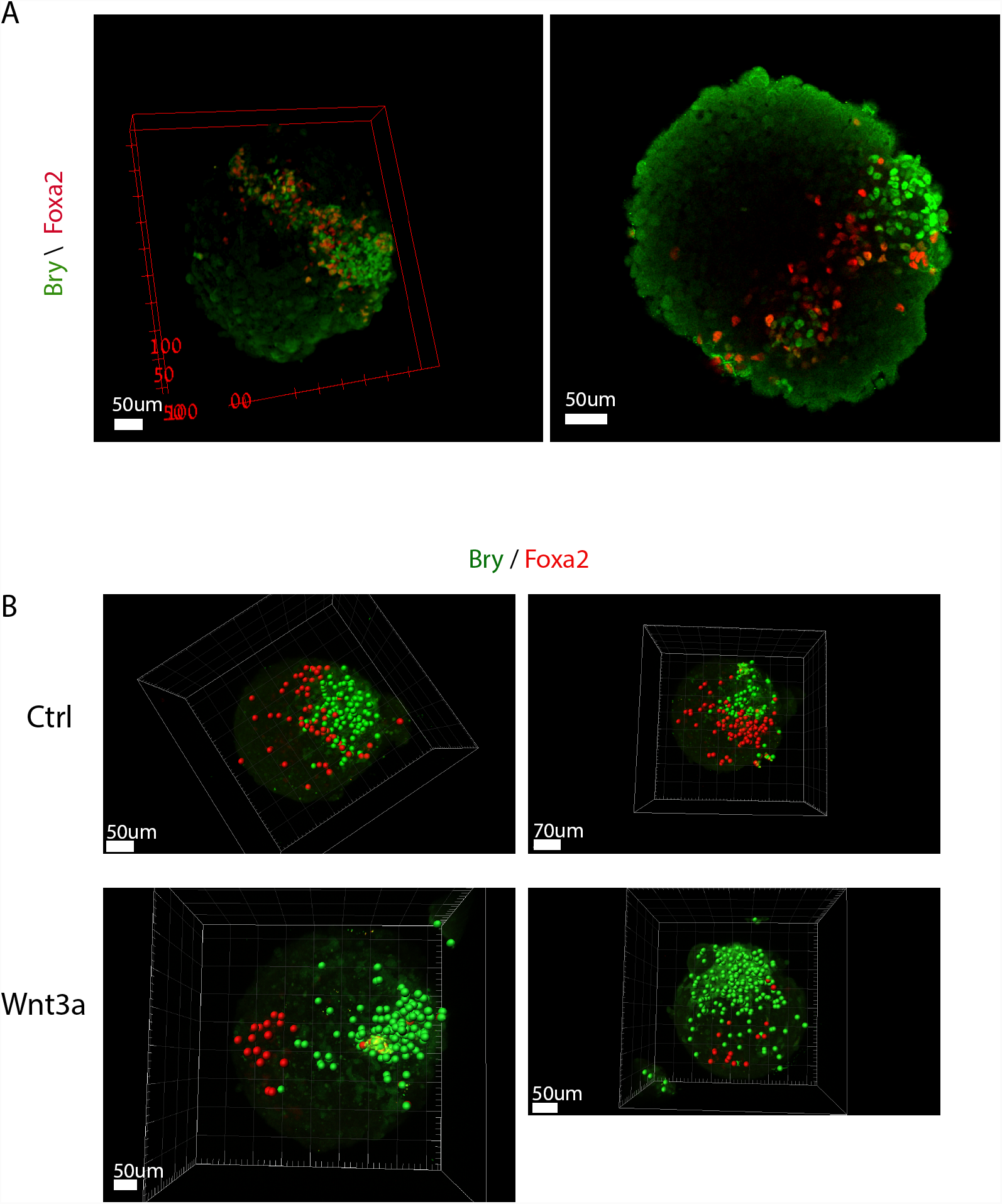
Bry and Foxa2 are spatially adjacent in EBs but can be decoupled by external Wnt signaling. (A) Immunostaining of Foxa2 and Bry at 72 hr from aggregation. Foxa2 spatially correlates with Bry expression (B) Segmentation of immunostained Foxa2 (red) and Bry (green) expressing cells at 72 hr from aggregation. Foxa2 is adjacent to Bry locus in wild type EBs, while Wnt3a treated EBs show Foxa2 downregulation and spatial decoupling from Bry.

**Fig S4.**
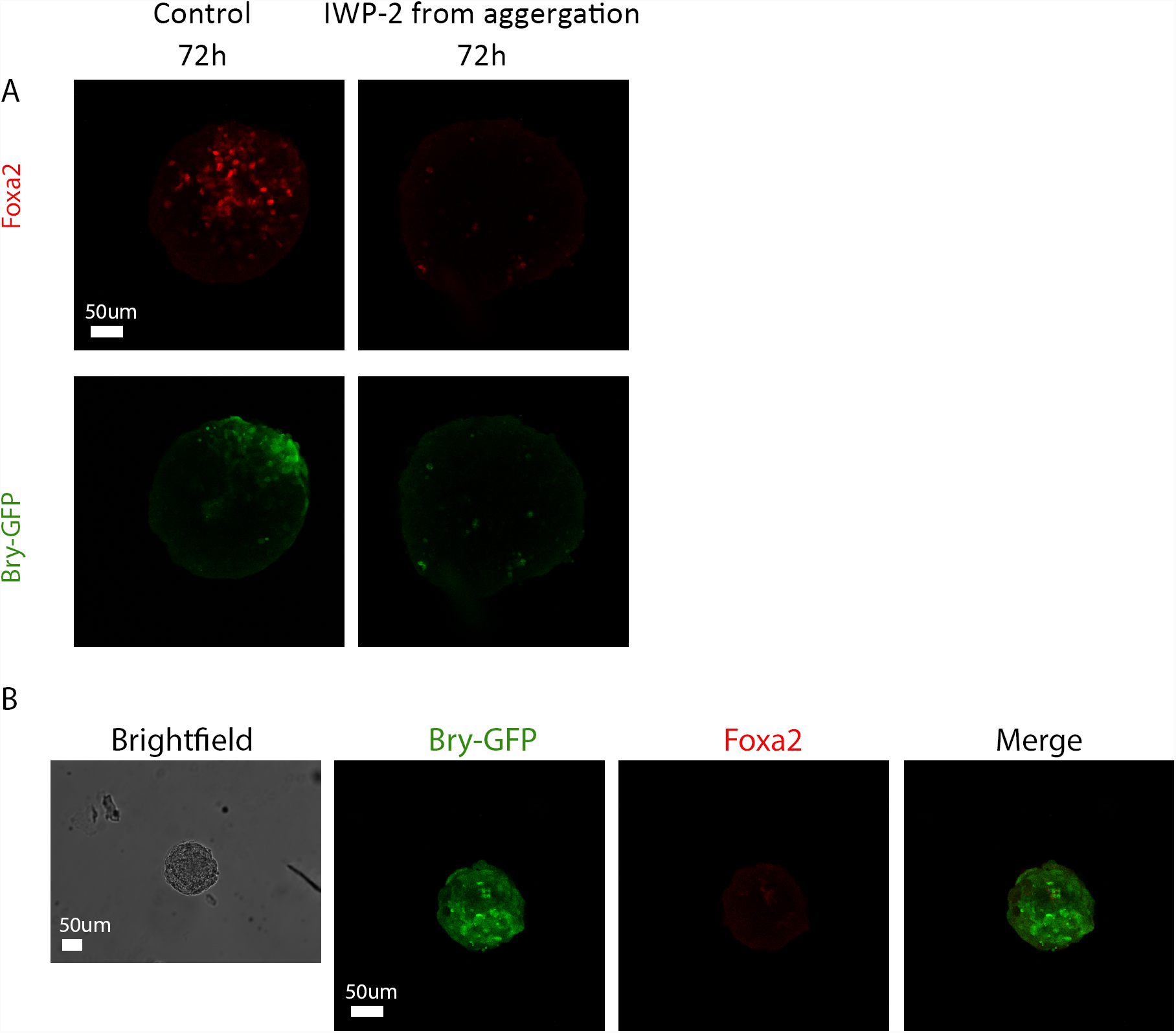
Foxa2 expression abolished under IWP-2 treatment or in small sized EBs. (A) Bry and Foxa2 are not expressed at 72 hr under IWP-2 perturbation starting at 0 hr from aggregation. (B) Small size EBs did not express Foxa2 at 72 hr.

**Movie S1, S2**

Three-dimensional time-lapse imaging of Bry-GFP in two E14 Bry-GFP embryoid bodies, imaged in microwells between 60 and 96 hr from aggregation and transfer to differentiation medium. Bry-GFP expression onsets at the bottom (contact point with the glass), and expanding upwards on the outer shell.

**Movie S3**

Epifluorescence time-lapse of a Bry-GFP embryoid body, where Bry onset occurred at the bottom. The EB is imaged between 24 and 90 hr from aggregation, where at 24 hr it was transferred to a microwell. Top row: left - brightfield imaging of the EB with its encompassing perimeter; center - Bry-GFP; right – overlay of brightfield and GFP-positive pixels (magenta). Bottom row: left - total Bry-GFP+ pixels vs. time point (time interval between points – 30 minutes). Blue line - raw data; red dashed line: alpha filter smoothing; yellow line: Bry onset threshold defined as 500 GFP+ pixels; center – EB radius vs. timepoint. Noise at higher time points is due to manual estimation of radius; right – snapshot of Bry-GFP onset frame.

**Movie S4**

Similar to Movie S3, for a case where Bry expression onset occurred at the contact point with the microwell side. Note in this case the EB originated from 3 smaller EBs that merged together after transfer to the microwell.

**Movie S5**

Similar to Movie S3, for an EB differentiated in an elongated micropool (width=200um). In this case, Bry-GFP expression onsets from two different loci, at the two sides compressed against the well walls.

**Movie S6**

Similar to Movie S5, for a case where Bry expression onsets from one of the compressed sides of the EB.

**Movie S7**

Time lapse of small EBs differentiated while embedded in Matrigel. Bry onset occurs uniformly from the whole sphere.

**Movie S8**

A Bry-GFP, pCMV-Strawberry large EB differentiated while embedded in Matrigel. Bry expression onsets from one locus, expanding from that point into the whole sphere.

